# Time without clocks: Human time perception based on perceptual classification

**DOI:** 10.1101/172387

**Authors:** Warrick Roseboom, Zafeirios Fountas, Kyriacos Nikiforou, David Bhowmik, Murray Shanahan, Anil K. Seth

## Abstract

Despite being a fundamental dimension of experience, how the human brain generates the perception of time remains unknown. Here, we provide a novel explanation for how human time perception might be accomplished, based on non-temporal perceptual classification processes. To demonstrate this proposal, we built an artificial neural system centred on a feed-forward image classification network, functionally similar to human visual processing. In this system, input videos of natural scenes drive changes in network activation, and accumulation of salient changes in activation are used to estimate duration. Estimates produced by this system match human reports made about the same videos, replicating key qualitative biases, including differentiating between scenes of walking around a busy city or sitting in a cafe or office. Our approach provides a working model of duration perception from stimulus to estimation and presents a new direction for examining the foundations of this central aspect of human experience.

---

In recent decades, predominant models of human time perception have been based on the presumed existence of neural processes that continually track physical time-so called pacemakers-similar to the system clock of a computer ^1,2,3^. Clear neural evidence for pacemakers at psychologically-relevant timescales has not been forthcoming and so alternative approaches have been suggested (e.g. ^4,5,6,7^). The leading alternative proposal is the network-state dependent model of time perception, which proposes that time is tracked by the natural temporal dynamics of neural processing within any given network ^8,9,10^. While recent work suggests that network-state dependent models may be suitable for describing temporal processing on short time scales ^11,12,10^, such as may be applicable in motor systems ^9,12,13,14^, it remains unclear how this approach might accommodate longer intervals (*>* 1s) associated with subjective duration estimation.

In proposing neural processes that attempt to track physical time as the basis of human subjective time perception, both the pacemaker and state-dependent network approaches stand in contrast with the classical view in both philosophical ^15^ and behavioural work ^16,17,18^ on time perception that emphasises the key role of perceptual content, and most importantly *changes* in perceptual content, to subjective time. It has often been noted that human time perception is characterised by its many deviations from veridical perception ^19,20,21,22^. These observations pose substantial challenges for models of subjective time perception that assume subjective time attempts to track physical time precisely. One of the main causes of deviation from veridicality lies in basic stimulus properties. Many studies have demonstrated the influence of stimulus characteristics such as complexity ^16,23^ and rate of change ^24,25,26^ on subjective time perception, and early models in cognitive psychology emphasised these features ^27,16,28,29^. Subjective duration is also known to be modulated by attentional allocation to the tracking time (e.g. prospective versus retrospective time judgements ^30,31,32,33,34^ and the influence of cognitive load ^35,32,33^).

Attempts to integrate a content-based influence on time perception with pacemaker accounts have hypothesized spontaneous changes in clock rate (e.g. ^36^), or attention-based modulation of the efficacy of pacemakers ^30,37^, while no explicit efforts have been made to demonstrate the efficacy of state-dependent network models in dealing with these issues. Focusing on pacemaker-based accounts, assuming that content-based differences in subjective time are produced by attention-related changes in pacemaker rate or efficacy implies a specific sequence of processes and effects. Firstly, it is necessary to assume that content alters how time is tracked, and that these changes cause pacemaker/accumulation to deviate from veridical operation. Changed pacemaker operation then leads to altered reports of time specific to that content. In contrast to this approach, we propose that the intermediate step of a modulated pacemaker, and the pacemaker in general, be abandoned altogether. Instead, we propose that *changes in perceptual content* can be tracked directly in order to determine subjective time. A challenge for this proposal is that it is not immediately clear how to quantify perceptual change in the context of natural ongoing perception. However, recent progress in machine learning provides a solution to this problem.

Accumulating evidence supports both the functional and architectural similarities of deep convolutional image classification networks (e.g. ^38^) to the human visual processing hierarchy ^39,40,41,42^. Changes in perceptual content in these networks can be quantified as the collective difference in activation of neurons in the network to successive inputs, such as consecutive frames of a video. We propose that this simple metric provides a sufficient basis for subjective time estimation. Further, because this metric is based on perceptual classification processes, we hypothesise that the produced duration estimates will exhibit the same content-related biases as characterise human time perception. To test our proposal, we implemented a model of time perception using an image classification network ^38^ as its core, and compared its performance to that of human participants in estimating time for the same natural video stimuli.

## Results

The stimuli for human and machine experiments were videos of natural scenes, such as walking through a city or the countryside, or sitting in an office or cafe (see Supplementary Video 1; Fig. 1D). These videos were split into durations between 1 and 64 seconds and used as the input from which our model would produce estimates of duration (see Methods for more details). To validate the performance of our model, we had human participants watch these same videos and make estimates of duration using a visual analogue scale (Fig. 1). Participants’ gaze position was also recorded using eye-tracking while they viewed the videos.

**Figure 1:**
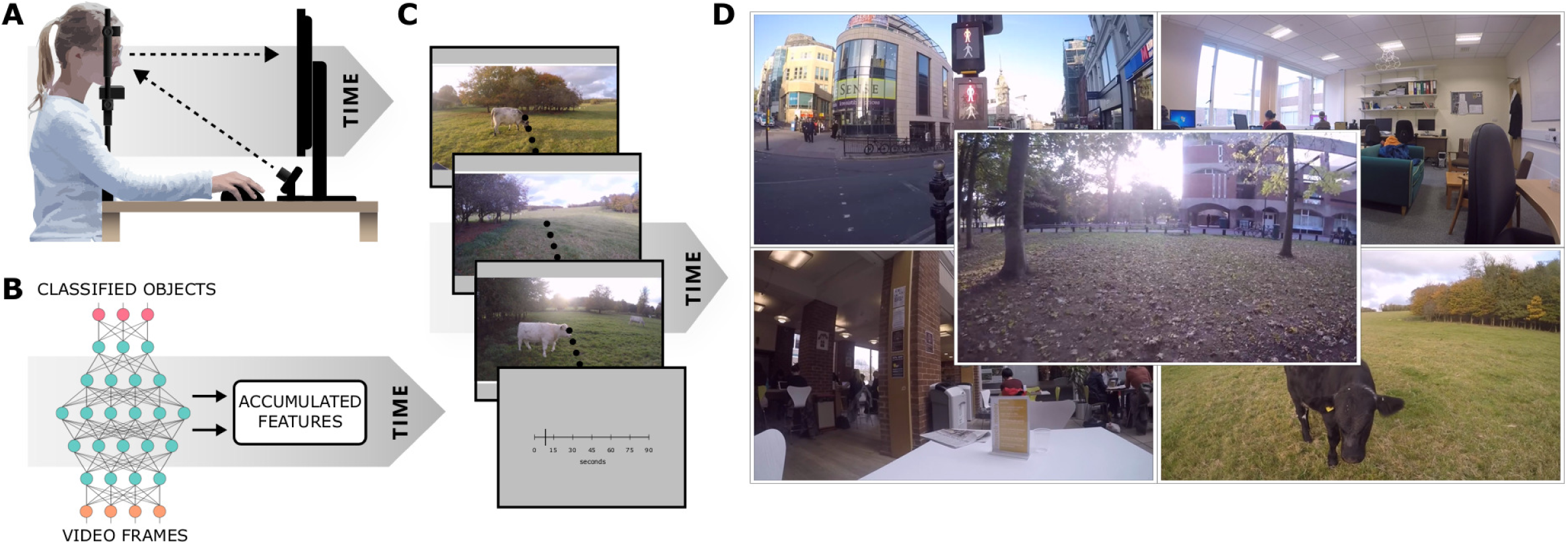
Experimental apparatus and procedure. (A) Human participants observed videos of natural scenes and reported the apparent duration while we tracked their gaze direction. (B) Depiction of the high-level architecture of the system used for simulations (Fig. 2). (C) Frames from a video used as a stimulus for human participants and input for simulated experiments. Human participants provided reports of the duration of a video in seconds using a visual analogue scale. (D) Videos used as stimuli for the human experiment and input for the system experiments included scenes recorded walking around a city (top left), in an office (top right), in a cafe (bottom left), walking in the countryside (bottom right) and walking around a leafy campus (centre).

The videos were input to a pre-trained feed-forward image classification network ^38^. To estimate time, the system measured whether the Euclidean distance between successive activation patterns within a given layer, driven by the video input, exceeded a dynamic threshold (Fig. 2). The dynamic threshold was implemented for each layer, following a decaying exponential corrupted by Gaussian noise and resetting whenever the measured Euclidean distance exceeded it. For a given layer, when the activation difference exceeded the threshold a salient perceptual change was determined to have occurred, and a unit of subjective time was accumulated (see Supplemental Results for model performance under a static threshold). To transform the accumulated, abstract temporal units extracted by the system into a measure of time in standard units (seconds) for comparison with human reports, we trained a Support Vector Machine (SVM) to estimate the duration of the videos based on the accumulated salient changes across network layers. Importantly, the regression was trained on the *physical* durations of the videos, not the human provided estimates. Therefore an observed correspondence between system and human-produced estimates would demonstrate the ability of the underlying perceptual change detection and accumulation method to model human duration perception, rather than the more trivial task of mapping human reports to specific videos/durations (see Methods for full details of system design and training).

**Figure 2:**
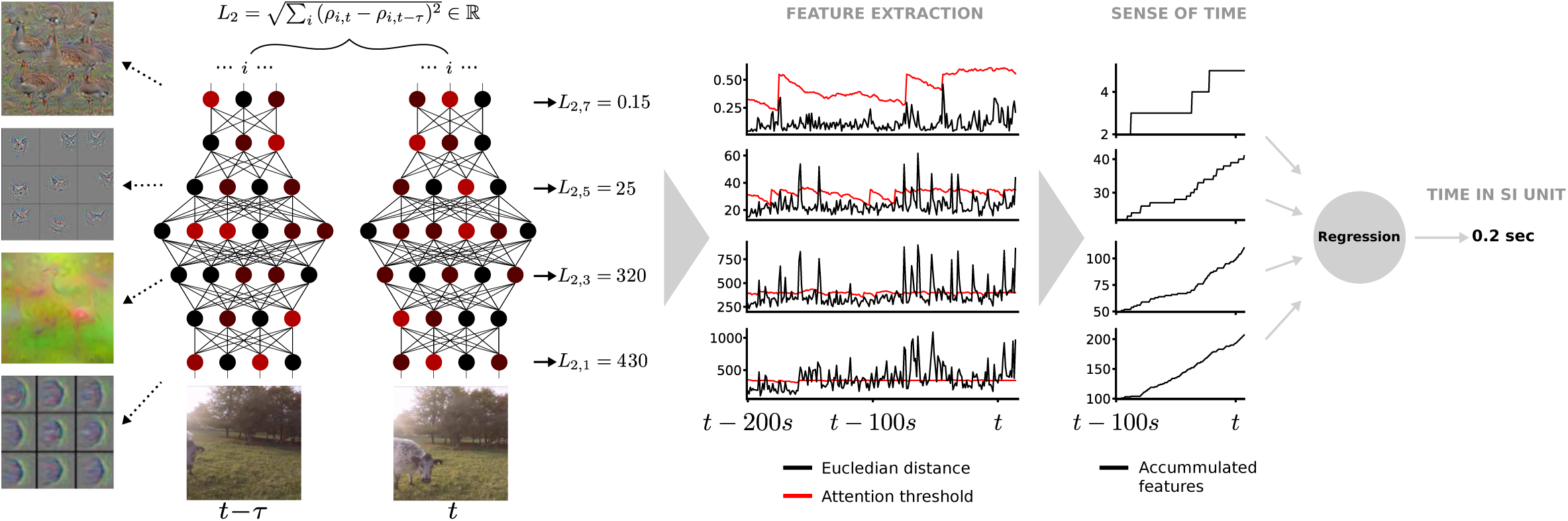
Depiction of the time estimation system. Salient changes in network activation driven by video input are accumulated and transformed into standard units for comparison with human reports. The left side shows visualizations of archetypal features to which layers in the classification network are responsive (adapted from ^43,44,45^). The bottom left shows two consecutive frames of video input. The connected coloured nodes depict network structure and activation patterns in each layer in the classification network for the inputs. *L*_2_ gives the Euclidean distance between network activations to successive inputs for a given network layer (layers conv2, pool5, fc7, output). In the Feature Extraction stage, the value of *L*_2_ for a given network layer is compared to a dynamic threshold (red line). When *L*_2_ exceeds the threshold level, a salient perceptual change is determined to have occurred, a unit of subjective time is determined to have passed, and is accumulated to form the base estimate of time. A regression method (support vector regression) is applied to convert this abstract time estimate into standard units (seconds) for comparison with human reports.

### Tracking changes in perceptual classification produces human-like time estimation

We initially had the system produce estimates under two input scenarios. In one scenario, the entire video frame was used as input to the network. In the other, input was spatially constrained by biologically relevant filtering-the approximation of human visual spatial attention by a ‘spotlight’ centred on real human gaze fixation. The extent of this spotlight approximated an area equivalent to human parafoveal vision and was centred on the participants’ fixation measured for each time-point in the video. Only the pixels of the video inside this spotlight were used as input to the system (see Supplementary Video 2).

As time estimates generated by the system were made on the same videos as the reports made by humans, human and system estimates could be compared directly. Fig. 3A shows duration estimates produced by human participants and the system under the different input scenarios. Participants’ reports demonstrated qualities typically found for human estimates of time: overestimation of short durations and underestimation of long durations (regression of responses to the mean/Vierordt’s law), and variance of reports proportional to the reported duration (scalar variability/Weber’s law). System estimates produced when the full video frame was input (Fig. 3B; Full-frame model) revealed qualitative properties similar to human reports-though the degree of over and underestimation was exaggerated, the variance of estimates were generally proportional to the estimated duration. These results demonstrate that the basic method of our system-accumulation of salient changes in activation of a perceptual classification network-can produce meaningful estimates of time. Specifically, the slope of estimation is non-zero with short durations discriminated from long, and the estimates replicate qualitative aspects of human reports often associated with time perception (Vierordt’s law and scalar variability). However, the overall performance of the system under these conditions still departed from that of human participants (Fig. 3E, F). (see Supplemental Results for results of experiments conducted on pixel-wise differences in the raw video alone, by-passing network activation).

**Figure 3:**
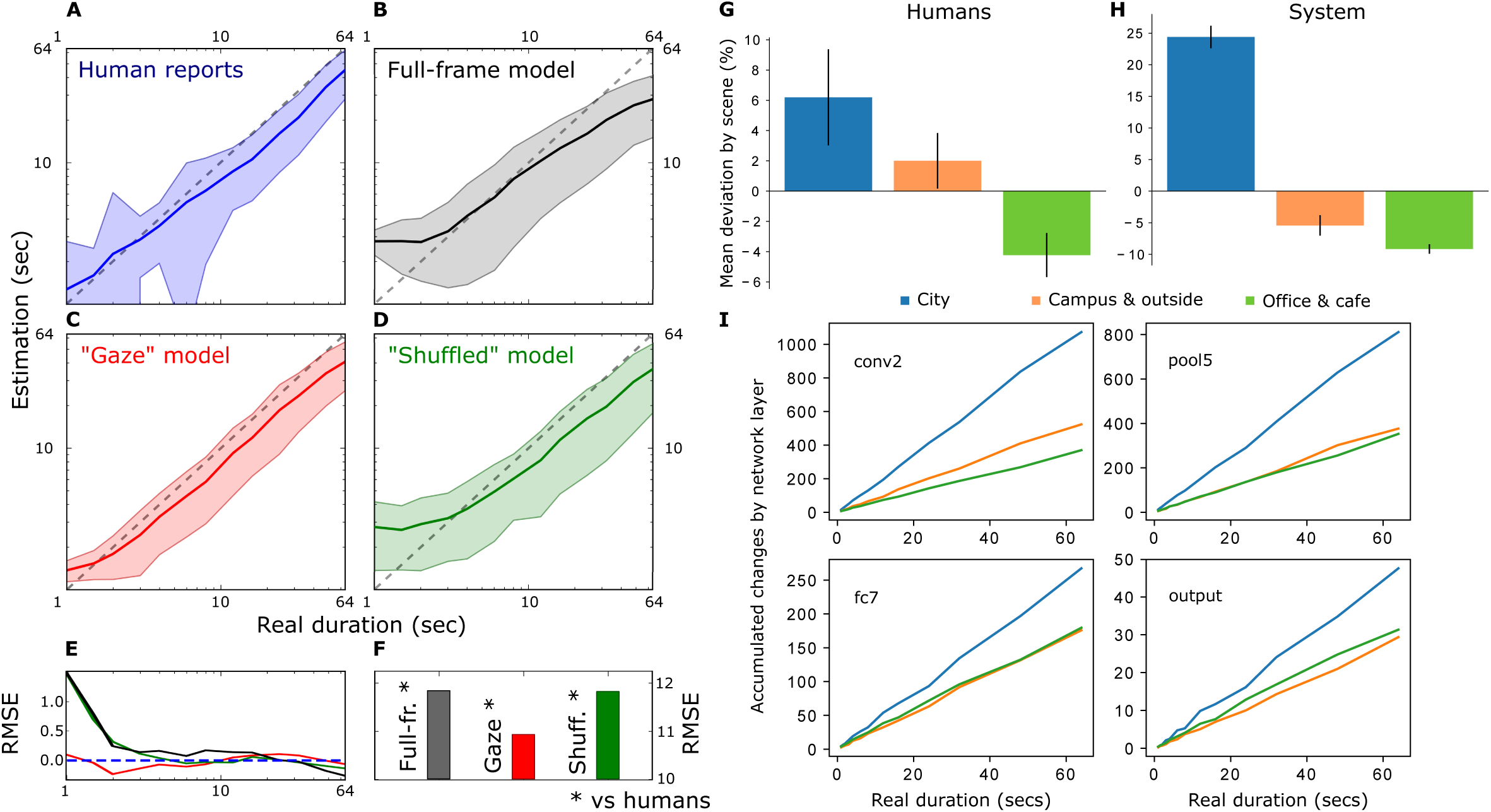
The mean duration estimates for 4290 trials for both human (A) and system (B,C,D) for the range of presented durations (1-64s). Shaded areas show *±*1 standard deviation of the mean. Human reports (A) show typical qualities of human temporal estimation with overestimation of short and underestimation of long durations. (B) System estimates when input the full video frame replicate similar qualitative properties, but temporal estimation is poorer than humans. (C) System estimates produced when the input was constrained to approximate human visual-spatial attention, based on human gaze data, very closely approximated human reports made on the same videos. When the gaze contingency was “Shuffled” such that the spotlight was applied to a different video than that from which it was obtained (D), performance decreases. (E) Comparison of mean absolute error between different estimations across presented durations. (F) Comparison of the root mean squared error of the system estimates compared to the human data. The “Gaze” model is most closely matched. (G) Mean deviation of duration reports by scene type, relative to mean duration estimation for human participants (mean shown in A). (H) Mean deviation of duration estimations by scene type, relative to mean duration estimation for the “Gaze” model (mean shown in C). (I) The number of accumulated salient perceptual changes over time in the different network layers (lowest to highest: conv2, pool5, fc7, output), depending on input scene type, for the “Gaze” model shown in (H). Error bars in (G) and (H) show standard error of the mean.

### Human-like gaze improves model performance

When the video input to the system was constrained to approximate human visual spatial attention by taking into account gaze position (“Gaze” model; Fig. 3C), system-produced estimates more closely approximated reports made by human participants (Fig. 3C, E, F), with substantially improved estimation as compared to estimates based on the full frame input. This result was not simply due to the spatial reduction of input caused by the gaze-contingent spatial filtering, nor the movement of the input frame itself, as when the gaze-contingent filtering was applied to videos other than the one from which gaze was recorded (i.e. gaze recorded while viewing one video then applied to a different video; “Shuffled” model), system estimates were poorer (Fig. 3D). These results indicate that the contents of where humans look in a scene play a key role in time perception and indicate that our approach is capturing key features of human time perception, as model performance is improved when input is constrained to be more human-like.

### Model and human time estimation vary by content

As described in the introduction, human estimates of duration are known to vary by content (e.g. ^16,23,24,25,26^). In our test videos, three different scenes could be broadly identified: scenes filmed moving around a city, moving around a leafy university campus and surrounding countryside, or from relatively stationary viewpoints inside a cafe or office (Fig. 1D). We reasoned that busy scenes, such as moving around a city, would generally provide more varied perceptual content, with content also being more complex and changing at a faster rate during video presentation. This should mean that city scenes would be judged as longer relative to country/outside and office or cafe scenes. As shown in (Fig. 3G), the pattern of biases in human reports is consistent with this hypothesis. Compared to the global mean estimates (Fig. 3A), reports made about city scenes were judged to be approximately 6% longer than the mean, while more stationary scenes, such as in a cafe or office, were judged to be approximately 4% shorter than the overall mean estimation (See Supplemental Results for full human and model produced estimates for each tested duration and scene).

To test whether the system-produced estimates exhibited the same content-based biases seen in human duration reports, we examined how system estimates differed by scene type. Following the same reasoning as for the human data, busy city scenes should provide a more varied input, which should lead to more varied activation within the network layers, therefore greater accumulation of salient changes and a corresponding bias towards overestimation of duration. As shown in (Fig. 3H), when the system was shown city scenes, estimates were biased to be longer (˜24%) than the overall mean estimation, while estimates for country/outside (˜4%) or office/cafe (˜7%) scenes were shorter than average. The level of overestimation for city scenes was substantially larger than that found for human reports, but the overall pattern of biases was the same: city *>* campus/outside *>* cafe/office (^1^ see General Discussion for discussion of system redundancy and its impact on overestimation). It is important to note again here that the model estimation was not optimised to human data in any way. The support-vector regression method mapped accumulated perceptual changes across network layers to the *physical* durations of the videos. That the same pattern of biases in estimation is found indicates the power of the underlying method of accumulating salient changes in perceptual content to produce human-like time perception.

Looking into the system performance more deeply, it can be seen that the qualitative matches between human reports and model estimation do not simply arise in the final step of the architecture, wherein the state of the accumulators at each network layer is regressed against physical 7 duration using a support-vector regression scheme. Even in the absence of this final step, which transforms accumulated salient changes into 8 standard units of physical time (seconds), the system showed the same pattern of biases in accumulation for most durations, at most of the 9 examined network layers. More perceptual changes were accumulated for city scenes than for either of the other scene types, particularly in the 0 lower network layers (conv2 and pool5). Therefore, the regression technique used to transform the tracking of salient perceptual changes is not critical to reproduce these scene-wise biases in duration estimation, and is needed only to compare system performance with human estimation in commensurate units.^2^ While regression of accumulated network activation differences into standard units is not critical to reproducing human-like biases in duration perception, basing duration estimation in network activation is key to model performance. When estimates are instead derived directly from differences between successive frames (on a pixel-wise basis) of the video stimuli, by-passing the classification network entirely, generated estimates are substantially worse, and most importantly, do not closely follow human biases in estimation. See Supplemental Results section *Changes in classification network activation, not just stimulation, are critical to human-like time estimation* for more details.

### Accounting for the role of attention in time perception

The role of attention in human time perception has been extensively studied (see^32^ for review). One key finding is that when attention is not specifically directed to tracking time (retrospective time judgements), or actively constrained by other task demands (e.g. high cognitive load), duration estimates differ from when attention is, or can be, freely directed towards time ^35,31,33,32^. Our model is based on detection of salient changes in neural activation underlying perceptual classification. To determine whether a given change is salient, the difference between previous and current network activation is compared to a running threshold, the level of which can be considered to be attention to changes in perceptual classification – effectively attention to time in our conception.

Regarding the influence of the threshold on duration estimation, in our proposal the role of attention to time is intuitive: when the threshold value is high (the red line in Feature Extraction in Fig. 2 is at a higher level in each layer of the network), a larger difference between successive activations is required in order for a given change to be deemed salient (when you aren’t paying attention to something, you are less likely to notice it changing, but large changes will still be noticed). Consequently, fewer changes in perceptual content are registered within a given epoch and, therefore, duration estimates are shorter. By contrast, when the threshold value is low, smaller differences are deemed salient and more changes are registered, producing generally longer duration estimates. Within our model, it is possible to modulate the level of attention to perceptual change using a single scaling factor referred to as Attention Modulation (see description of Equation 1). Changing this scaling factor alters the threshold level which, following the above description, modulates attention to change in perceptual classification. Shown in Fig. 4B are duration estimates for the “Gaze” model presented in Fig. 3C under different levels of attentional modulation. Lower than normal attention levels lead to general underestimation of duration (lighter lines), while higher levels of attention lead to increasing overestimation (darker lines; see Supplemental Results for results modulating attention in the other models). Taking duration estimates produced under a lower level of attention and comparing with those produced under a higher level can produce the same pattern of differences in estimation often associated attending (high attention to time; prospective/low cognitive load) or not attending (low attention to time; prospective/high cognitive load) reported in the literature. These results demonstrate the flexibility of our model to deal with different demands posed from both “bottom-up” basic stimulus properties as well as “top-down” demands of allocating attention to time.

**Figure 4:**
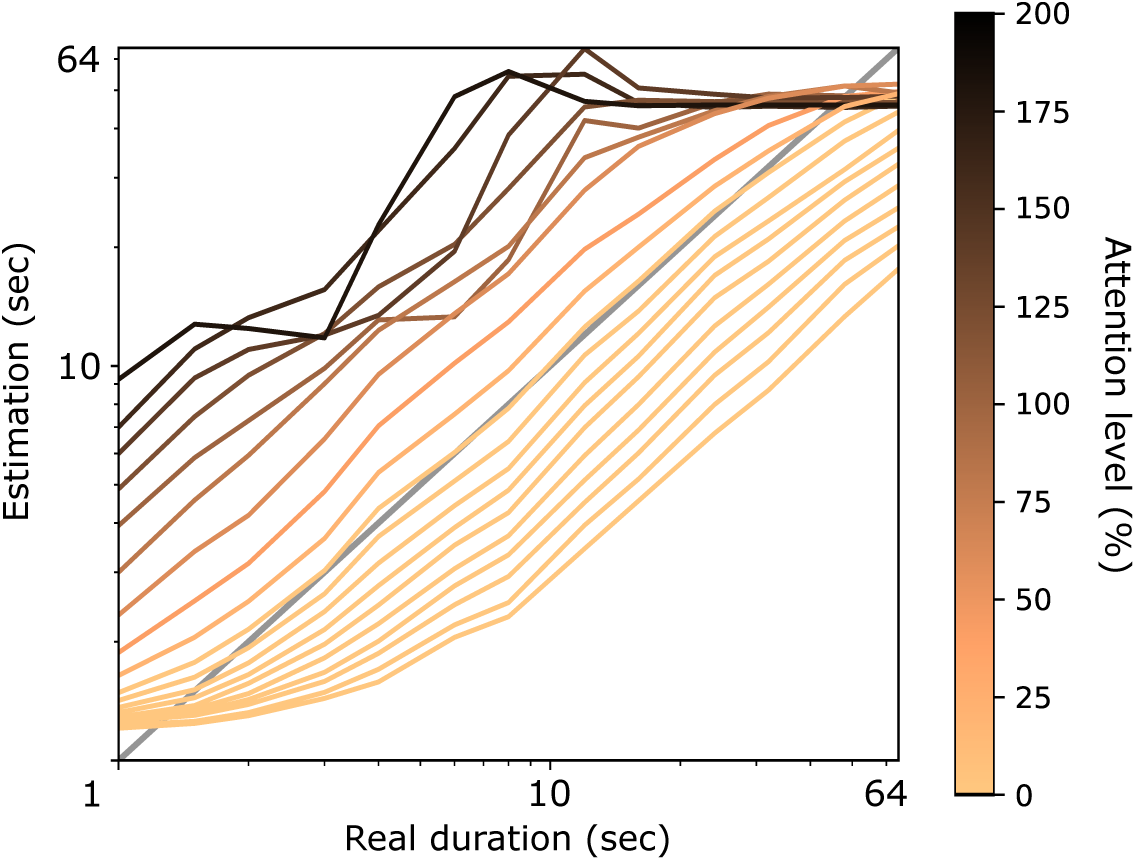
Comparison of system duration estimation at different levels of attentional modulation. Attentional modulation refers to a scaling factor applied to the parameters *T*_*max*_ and *T*_*min*_, specified in Table 1 and Equation 1. Changing the Attention level affects duration estimates, biasing estimation across a broad range of levels. The model still generally differentiates longer from shorter durations, as indicated by the positive slopes with increasing real duration, but also exhibits biases consistent with those known from behavioural literature associated with attention to time (e.g. ^33,32^).

**Figure 5:**
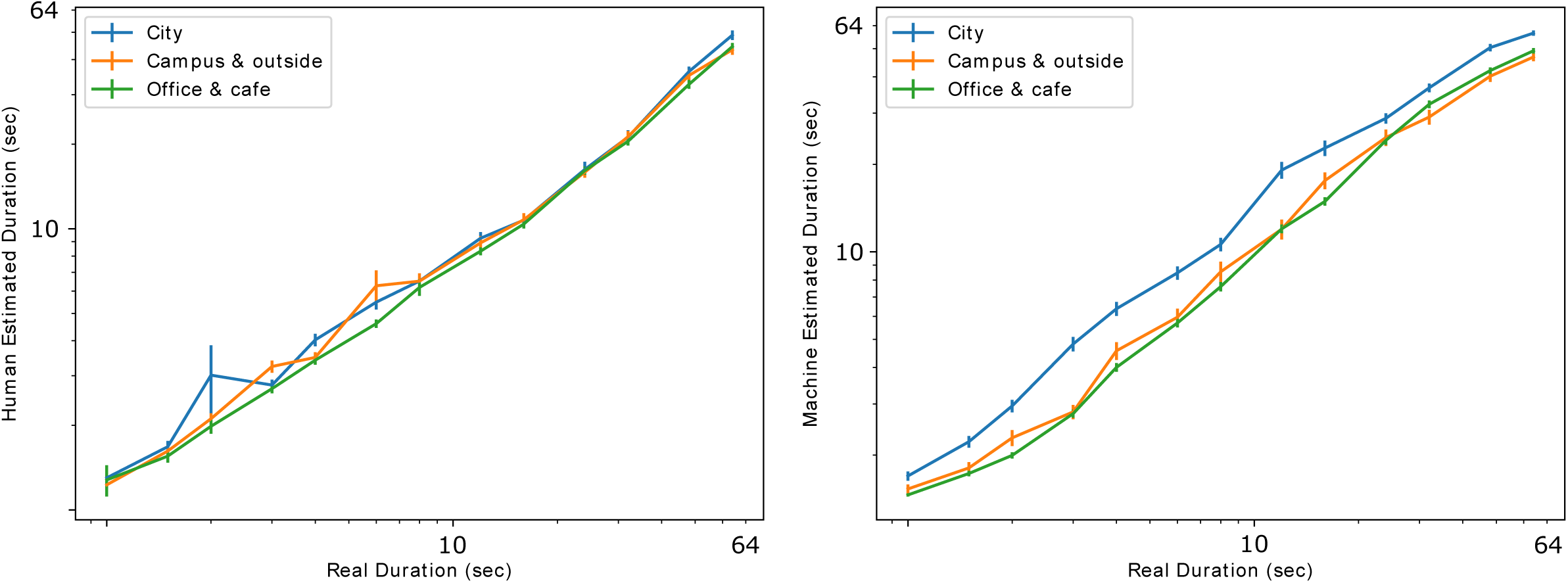
**(Supplementary figure)** Duration estimation for each tested video duration, separated by scene-type, for human (left panel) and model (right panel) experiments. Error bars indicate standard error of the mean. As shown in Fig. 3G and H in the main text, for both humans and the system, city scenes are typically estimated as longer than campus and outside, or office and cafe scenes. The degree of this overestimation is stronger for the system, but the overall pattern of results is the same for human and system estimation.

## Discussion

This study tested the proposal that accumulating salient changes in perceptual content, indicated by differences in successive activation of a perceptual classification network, would be sufficient to produce human-like estimates of duration. Results showed that system-produced estimates could differentiate short from long durations, supporting basic duration estimation. Moreover, when input to the system was constrained to follow human gaze, model estimation improved and became more like human reports. Model estimates were also found to vary by the kind of scene presented, producing the same pattern of biases in estimation seen in human reports for the same videos, with evidence for this bias present even within the accumulation process itself. Finally, we showed that within the proposed model, the ability to modulate the level of attention to perceptual changes produced systematic under-and overestimation of durations, consistent with the literature on the interaction of attention to time and duration estimation. Overall, these results provide compelling support for the hypothesis that human subjective time estimation can be achieved by tracking non-temporal perceptual classification processes, in the absence of any regular pacemaker-like processes.

One might worry that the reliance of our model on a visual classification network is a flaw; after all, it is clear that human time perception depends on more than vision alone, not least because blind people still perceive time. However, the proposal is for a simple conceptual and mechanistic basis to accomplish time perception under naturalistic conditions using complex stimuli. The model’s performance demonstrates the feasibility of this approach when benchmarked against human performance, revealing similar performance under similar constraints. It should be noted that the described human data was obtained with participants seated in a quiet room, and with no auditory track to the video. This created an environment in which the most salient changes during a trial were within the video presentation. Certainly, in an experiment containing audio, audition would contribute to reported duration – and in some cases move human performance away from that of our vision-only model. Similarly, if participants sat in a quiet room with no external stimulation presented, temporal estimations would likely be biased by changes in the internal bodily states of the observer. Indeed, the insula cortex has been suggested to track and accumulate changes in bodily states that contribute to subjective time perception ^46,47^.

Although the basis of our model is fundamentally visual-an image classification network-similar classification network models exist for audition (e.g. ^48,49^), suggesting the possibility to implement the same mechanism in models for auditory classification. This additional level of redundancy in estimation would likely improve performance for scenarios that include both visual and auditory information, as has been shown in other cases where redundant cues from different modalities are combined ^50,51,22,52,53^. Additional redundancy in estimation would also likely reduce the propensity for the model to overestimate in scenarios that contain many times more perceptual changes than expected (such as indicated by the difference between human and model scene-wise biases Fig. 3H and G). Inclusion of further modules such as memory for previous duration estimations is also likely to improve system estimation as it is now well-established that human estimation of duration depends not only on the current experience of a duration, but also past reports of duration ^54,55^ (see also below discussion of predictive coding). While future extensions of our model could incorporate modules dedicated to auditory, interoceptive, memory, and other processes, these possibilities do not detract from the fact that the current implementation provides a simple mechanism that can be applied to these many scenarios, and that when human reports are limited in a similar way to the model, human and model performance are strikingly similar.

The core conception of our proposal shares some similarities with the previously discussed state-dependent network models of time ^8,9^. As in our model, the state-dependent network approach suggests that changes in activation patterns within neural networks (network states) over time can be used to estimate time. However, rather than simply saying that any dynamic network has the capacity to represent time by virtue of its changing state, our proposal goes further to say that changes in perceptual classification networks are the basis of content-driven time perception. This position explicitly links time perception and content, and moves away from models of subjective time perception that attempt to track physical time, a notion that has long been identified as conceptually problematic ^19^. A primary feature of state-dependent network models is their natural opposition to the classic depiction of time perception as a centralised and unitary process ^56,11,57^, as suggested in typical pacemaker-based accounts ^1,2,3^. Our suggestion shares this notion of distributed processing, as the information regarding salient changes in perceptual content within a specific modality (vision in this study) is present locally to that modality.

Finally, the described model used Euclidean distance in network activation as the metric of difference between successive inputs-our proxy for perceptual change. While this simple metric was sufficient to deliver a close match between model and human performance, future extensions may consider alternative metrics. The increasingly influential a predictive coding approach to perception ^58,59,60,61,62^ suggests one such alternative which may increase the explanatory power of the model. Predictive coding accounts are based on the idea that perception is a function of both prediction and current sensory stimulation. Specifically, perceptual content is understood as the brain’s “best guess” (Bayesian posterior) of the causes of current sensory input, constrained by prior expectations or predictions. In contrast to bottom-up accounts of perception, in which perceptual content is determined by the hierarchical elaboration of afferent sensory signals, strong predictive coding accounts suggest that bottom-up signals (i.e., flowing from sensory surfaces inwards) carry only the prediction errors (the difference, at each layer in a hierarchy, between actual and predicted signals), with prediction updates passed back down the hierarchy (top-down signals) to inform future perception. A role for predictive coding in time perception has been suggested previously, both in specific ^6^ and general models ^63^, and as a general principle to explain behavioural findings ^64,65,66,67^. Our model exhibits the basic properties of a minimal predictive coding approach; the current network activation state is the best-guess (prediction) of the future activation state, and the Euclidean distance between successive activations is the prediction error. The good performance and robustness of our model may reflect this closeness in implementation. While our basic implementation already accounts for some context-based biases in duration estimation (e.g. scene-wise bias), future implementations can include more meaningful “top-down”, memory and context driven constraints on the predicted network state (priors) that will account for a broader range of biases in human estimation.

In summary, subjective time perception is fundamentally related to changes in perceptual content. Here we show that a system built upon detecting salient changes in perceptual content across a hierarchical perceptual classification network can produce human-like time perception for naturalistic stimuli. Critically, system-produced time estimates replicated well-known features of human reports of duration, with estimation differing based on biologically relevant cues, such as where in a scene attention is directed, as well as the general content of a scene (e.g. city or countryside, etc). Moreover, we demonstrated that modulation of the threshold mechanism used to detect salient changes in perceptual content provide the capacity to reproduce the influence of attention to time in duration estimation. That our system produces human-like time estimates based on only natural video inputs, without any appeal to a pacemaker or clock-like mechanism, represents a substantial advance in building artificial systems with human-like temporal cognition, and presents a fresh opportunity to understand human perception and experience of time.

## Acknowledgments

This work was supported by the European Union Future and Emerging Technologies grant (GA:641100) TIMESTORM – Mind and Time: Investigation of the Temporal Traits of Human-Machine Convergence and the Dr. Mortimer and Theresa Sackler Foundation, supporting the Sackler Centre for Consciousness Science. Thanks to Michaela Klimova, Francesca Simonelli and Virginia Mahieu for assistance with the human experiment. Thanks to Tom Wallis and Andy Philippides for comments on previous versions of the manuscript.

## Methods

### Participants

Participants were 55 adults (21.2 years, 40 female) recruited from the University of Sussex, participating for course credit or £5 per hour. Participants typically completed 80 trials in the 1 hour experimental session, though due to time or other constraints some participants only completed as few as 20 trials (see Supplemental Data for specific trial completion details). This experiment was approved by the University of Sussex ethics committee.

### Apparatus

Experiments were programmed using Psychtoolbox 3^71,72,73^ in MATLAB 2012b (MathWorks Inc., Natick, US-MA) and the Eyelink Toolbox ^74^, and displayed on a LaCie Electron 22 BLUE II 22” with screen resolution of 1280 x 1024 pixels and refresh rate of 60 Hz. Eye tracking was performed with Eyelink 1000 Plus (SR Research, Mississauga, Ontario, Canada) at a sampling rate of 1000 Hz, using a desktop camera mount. Head position was stabilized at 57 cm from the screen with a chin and forehead rest.

### Stimuli

Experimental stimuli were based on videos collected throughout the City of Brighton in the UK, the University of Sussex campus, and the local surrounding area. They were recorded using a GoPro Hero 4 at 60 Hz and 1920 x 1080 pixels, from face height. These videos were processed into candidate stimulus videos 165 minutes in total duration, at 30 Hz and 1280 x 720 pixels. To create individual trial videos, a psuedo-random list of 4290 trials was generated-330 repetitions of each of 13 durations (1, 1.5, 2, 3, 4, 6, 8, 12, 16, 24, 32, 48, 64s). The duration of each trial was psuedo-randomly assigned to the equivalent number of frames in the 165 minutes of video. There was no attempt to restrict overlap of frames between different trials. The complete trial list and associated videos are available in the Supplemental Data.

For computational experiments when we refer to the “Full Frame” we used the center 720 x 720 pixel patch from the video (56 percent of pixels; approximately equivalent to 18 degrees of visual angle (dva) for human observers). When computational experiments used human gaze data, a 400 x 400 pixel patch was centered on the gaze position measured from human participants on that specific trial (about 17 percent of the image; approximately 10 dva for human observers).

### Computational model architecture

The computational model is made up of four parts: 1) An image classification deep neural network, 2) a threshold mechanism, 3) a set of accumulators and 4) a regression scheme. We used the convolutional deep neural network AlexNet ^38^ available through the python library caffe ^75^. AlexNet had been pretrained to classify high-resolution images in the LSVRC-2010 ImageNet training set ^76^ into 1000 different classes, with state-of-the-art performance. It consisted of five convolutional layers, some of which were followed by normalisation and max-pooling layers, and two fully connected layers before the final 1000 class probability output. It has been argued that convolutional networks’ connectivity and functionality resemble the connectivity and processing taking place in human visual processing ^40^ and thus we use this network as the main visual processing system for our computational model. At each time-step (30 Hz), a video frame was fed into the input layer of the network and the subsequent higher layers were activated. For each frame, we extracted the activations of all neurons from layers conv2, pool5, fc7 and the output probabilities. For each layer, we calculated the Euclidean distance between successive states. If the activations were similar, the Euclidean distance would be low, while the distance between neural activations corresponding to frames which include different objects would be high.

A “temporal attention” mechanism was implemented to dynamically calibrate the detection of changes between neural activations (threshold) resulting from successive frames. Each network layer had an initial threshold value for the distance in neural space. This threshold decayed with some stochasticity (Eq. 1) over time, in order to replicate the role of normalisation of neural responses to stimulation over time ^77,78^. When the measured Euclidean distance in a layer exceeded the threshold, the counter in that layers’ accumulators was incremented by one and the threshold of that layer was reset to its maximum value. The purpose of a decaying function was to accommodate time perception across various environments with exceptionally few or exceptionally many features. However, time estimation was still possible with a static threshold (see Supplemental Results: Model performance does not depend on threshold decay). Implementation details for each layer can be found in the table below, and the threshold was calculated as:

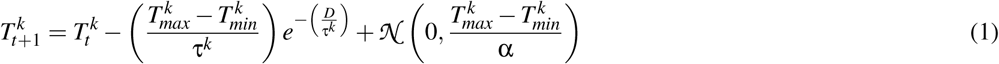

where 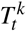 is the threshold value of *k*^*th*^ layer at timestep *t* and *D* indicates the number of timesteps since the last time the threshold value was reset. 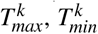 and τ^*k*^ are the maximum threshold value, minimum threshold value and decay timeconstant for *k*^*th*^ layer respectively, values which are provided in Table 1. Stochastic noise drawn from a Gaussian was added to the threshold and α-a dividing constant to adjust the variance of the noise. Finally, the level of attention was modulated by a global scaling factor *C >* 0 applied to the values of 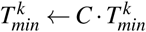 and 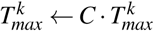.

**Table 1:** Threshold mechanism parameters

| Parameters for implementing salient event threshold |  |  |  |  |  |
| --- | --- | --- | --- | --- | --- |
| Layer | No. neurons | $T_{max}$ | $T_{min}$ | $\tau$ | $\alpha$ |
| conv2 | 290400 | 340 | 100 | 100 | 50 |
| pool5 | 9216 | 400 | 100 | 100 | 50 |
| fc7 | 4096 | 35 | 5 | 100 | 50 |
| output | 1000 | 0.55 | 0.15 | 100 | 50 |

## Supplementary Results

### Content, not model regularity drives time estimation

A potential criticism of the results in the main text would be that they simply reflect the operation of another type of pacemaker, in this case one that underlies the updating of perceptual content. As calculation of salient network activation changes in the model occurs at some defined frequency (the video was input to the system, and activation difference calculated, at 30 Hz in the above results), one might suspect that our system is simply mimicking a physical pacemaker, with the regular updates taking the role of, in the most trivial example, the movement of hands on a clock face. However, it is easy to demonstrate that the regularity of model operation is not the predominant feature in determining time estimates. If it were, duration estimates for the “Gaze” versus “Shuffled” models would be highly similar, as they contain the same input rate (30 Hz) and temporal features induced by movement of the gaze spotlight. This is clearly not the case (Fig. 3C and Fig. 3D in main text).

To thoroughly reject the concern that the regularity of the system update rate was the main determinant of time estimation in our system, we compared the salient changes accumulated by the system when inputting the “normal” videos at 30 Hz, with accumulated changes under three conditions: videos in which the frame rate was halved (skipped every second frame), videos in which some frames were skipped psuedo-randomly with a frequency of 20%, or videos input at 30Hz, but with the video frames presented in a shuffled order. The results showed that the manipulations of frame rate (skipping every second frame or 20% of frames) produced only small differences in accumulated changes over time compared to the normal input videos (Fig. 6). However, when the input rate was kept at 30 Hz, but the presentation order of the frames shuffled, thereby disrupting the flow of content in the video, the number of accumulated changes was very different (up to around 40 times *more* different from standard than either the halved or randomly skipped frame cases; see Fig. 6). These results underline that our system was producing temporal estimates based predominantly on the content of the scene, not the update rate of the system.

**Figure 6:**
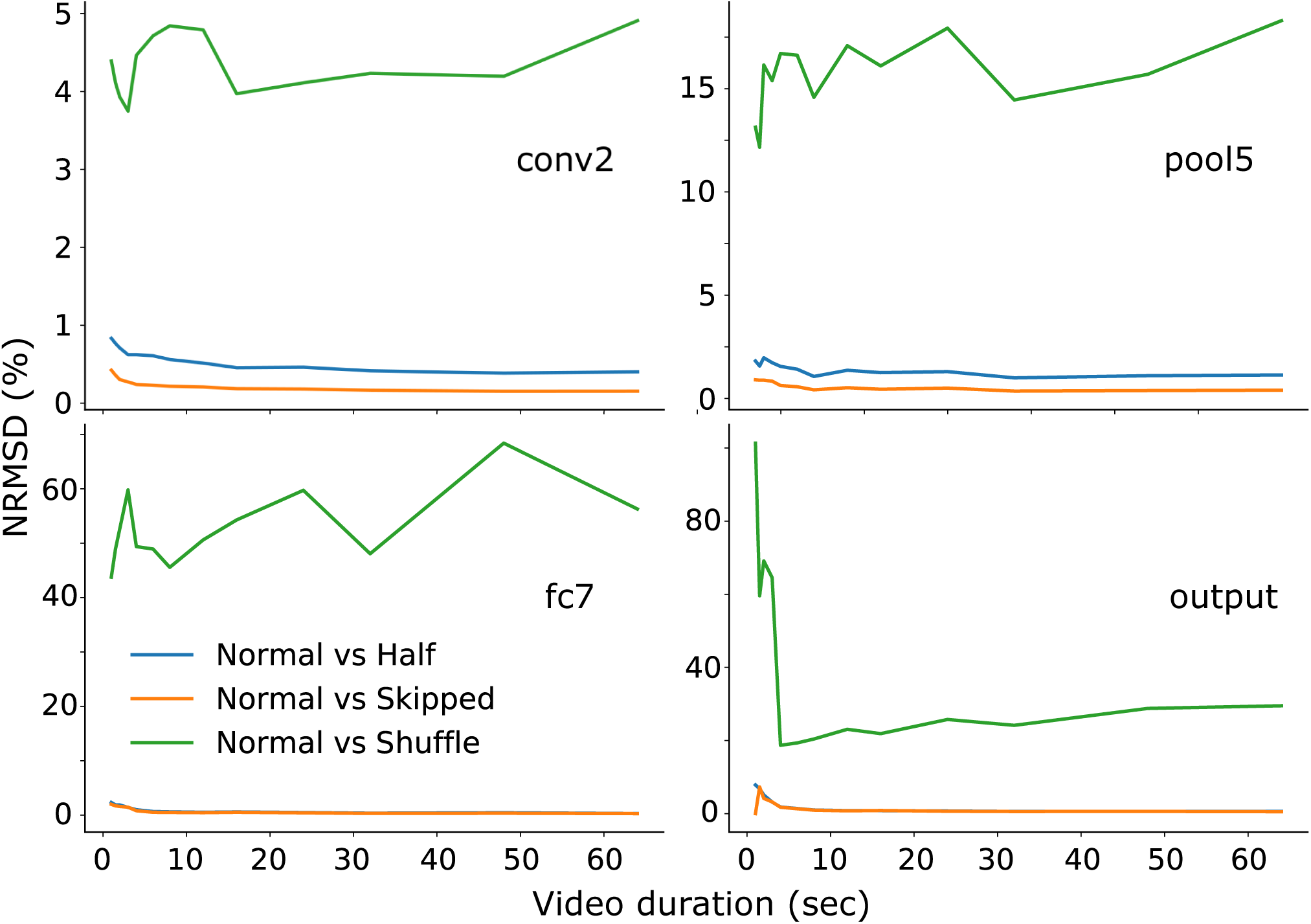
**(Supplementary figure)**(A) Comparison of system accumulation of salient changes depending on input frame rate and composition of the input video. Each panel shows the normalised root-mean squared difference between the accumulated salient changes in the system when given the normal input video at 30 Hz, compared to input videos at half the frame rate, inputs videos with 20% of frames pseudo-randomly skipped, and input videos presented at 30 Hz (same as the normal input videos), but with the order of presentation of the video frames shuffled. The manipulations of frame rate (halving or skipping 20%) had little effect on the accumulated changes (blue and orange lines), while shuffling the order of presentation of the frames altered the accumulation of salient changes dramatically (green lines).

### Model performance is robust across threshold parameters

The parameters of the model, 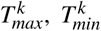 and τ^*k*^, were chosen so that the Euclidean distances for each layer exceeded the threshold only when a large increase occurred. The choice of particular values is not very important as the model performance is robust across a broad range of these values. When we scaled the values of 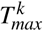 and 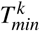 by a factor allowing us to vary the level of the threshold mechanism (*attention modulation*), our model could still estimate time with relatively good accuracy across a broad range of parameter values (Fig. 7Ai-Aiii) and, most importantly, still differentiate between short and long durations (slope is greater than zero for most levels). To further examine the effect of 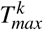 and 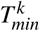, we scaled each parameter by an independent scaling factor to show that the model estimations (compared to the real physical duration) are robust over a wide range of values for these two parameters (Fig. 7B). These results show that system-produced estimation is relatively accurate (relative to physical duration) across a very broad range of parameters for 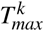 and 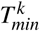.

**Figure 7:**
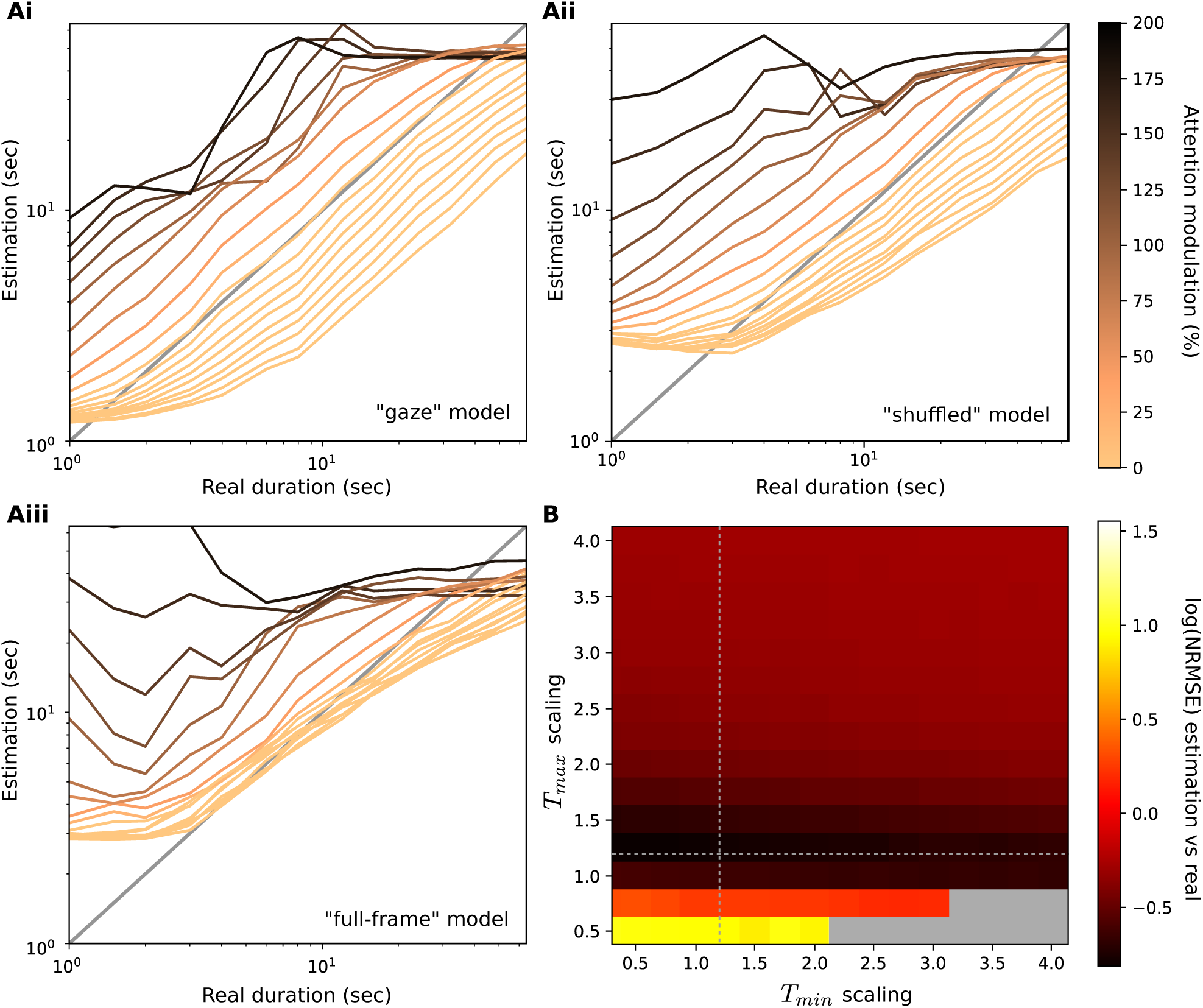
**(Supplementary figure)** Robustness of the temporal attention mechanism. **A:** Comparison of system duration estimation at different levels of attention modulation. This level refers to a scaling factor applied to the parameters *T*_*max*_ and *T*_*min*_, specified in Table 1. and Equation 1. Each panel shows the performance for a different variant of the model (“Gaze”, “Shuffled” and “Full-frame”). While changing the attention level did affect duration estimates, often resulting in a bias in estimation (e.g. many levels of the “Full-frame” exhibit a bias towards over-estimation; darker lines), across a broad range of Attention levels the models (particularly in the “Gaze” model) still differentiate longer from shorter durations, as indicated by the positive slopes with increasing real duration. For the models in Fig. 3, the following scalings were used: (“Gaze”: 1.20, “Shuffled”: 1.10 and “Full-frame”: 1.06) as they were found to produce estimations most closely matching human reports. **B:** Normalised root mean squared error (NRMSE) of duration estimations of the “Gaze” model versus real physical durations, for different combinations of values for the parameters *T*_*max*_ and *T*_*min*_ in Equation (1). The gray areas in the heatmap represent combinations of values that cannot be defined. Dotted lines represent the chosen attention threshold scaling used for the “Gaze” model in Fig. 3.

**Figure 8:**
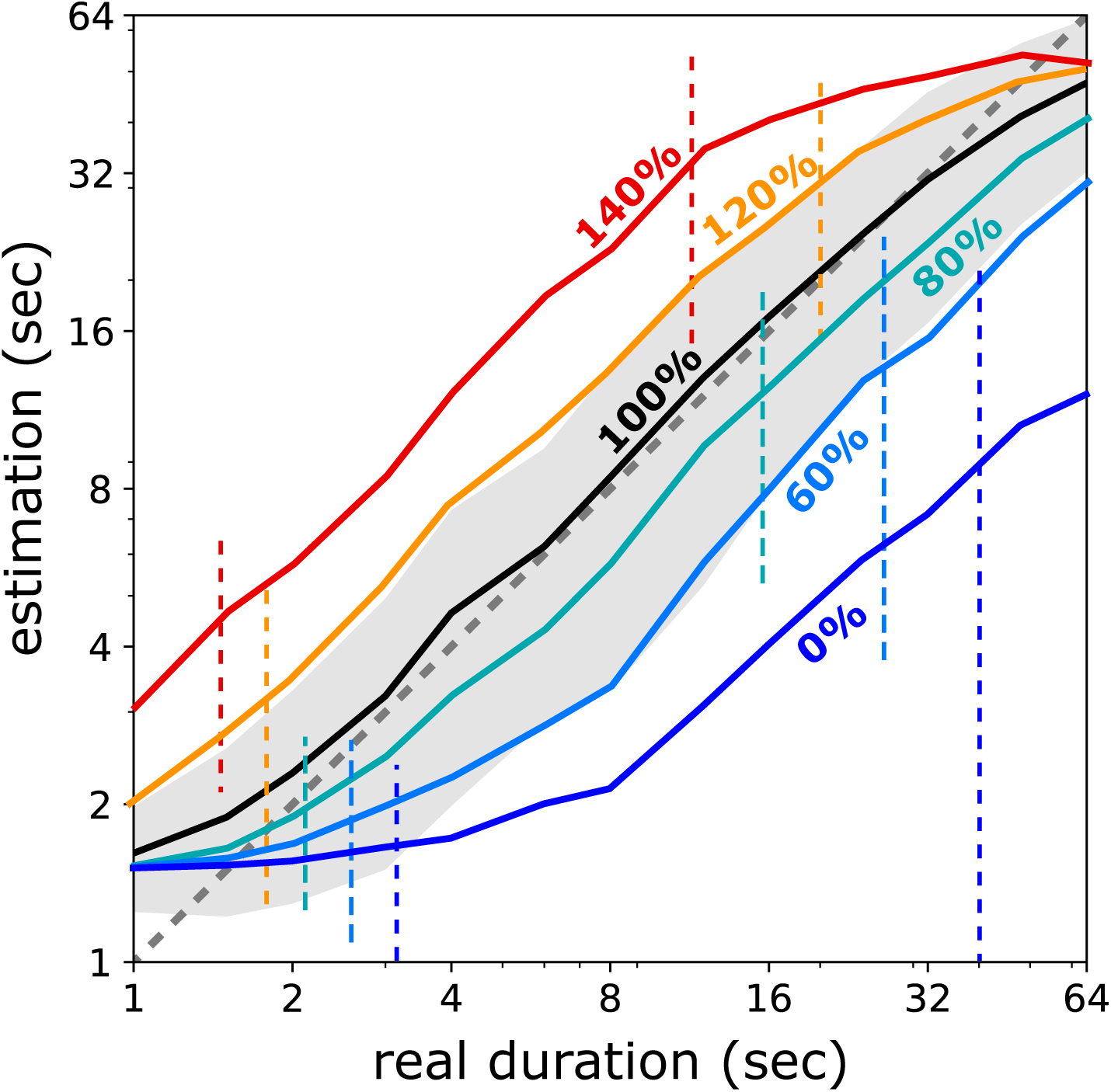
**(Supplementary figure)** Comparison of system estimation with fixed thresholds at different levels of attention modulation. As for estimation with dynamic thresholds (Fig. 7), the system can differentiate short from long durations effectively, and modulation of attention level causes a similar pattern of over and underestimation as found with the dynamic threshold.

### Model performance does not depend on threshold decay

The threshold used in the experiments reported in the main text included a noisy decay that continued until the threshold was exceeded, according to the parameters described in Equation 1. This decay was included to approximate the role of normalisation of neural response that is known to occur within the sensory systems (e.g. visual processing) the function of which we are attempting to mimic, and further, to facilitate model performance across a wider array of possible scene types and content. However, this decay is not essential for the model to perform well, discriminate short from long durations, and have the potential for attentional modulation. This can be seen if the threshold is set at a single level for all scenes and the regression mechanism is trained on accumulated salient changes under this single threshold level. As shown in Fig. 7, if the threshold is simply fixed as

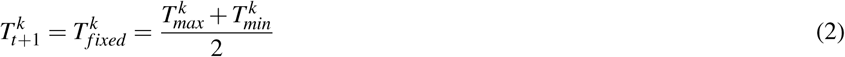

then the estimation remains similar to that reported in the main text (e.g. Fig. 3B-C). Furthermore, as discussed in the section *Accounting for the role of attention in time perception* in the main text, if this threshold level is changed by modulation of a global scaling factor *C >* 0 of *attention modulation*, system duration estimates become biased. In this case, the impact on the threshold when modulating attention canbe seen as 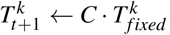, thus altering the probability that a given change between consecutive frames will be determined to be salient and accumulated to drive an increase in subjective duration. As a result, estimations become biased towards shorter estimations with a lower attention modulation, and longer estimations with higher attention modulation-consistent with the proposed interaction of attention and duration estimation covered in the main text. This effect shows that the dynamic nature of the threshold in the main implementation is not strictly necessary for meaningful estimates of time to be generated when tracking salient changes in network activation, and for those estimates to be modulated by attention to time.

### Model performance is not due to regression overfitting

The number of accumulated salient perceptual changes recorded in the accumulators represent the elapsed duration between two points in time. In order to convert estimates of subjective time into units of time in seconds, a simple regression method was used based on epsilon-Support Vector Regression (SVR) from sklearn python toolkit ^79^. The kernel used was the radial basis function with a kernel coefficient of 10^*-*4^ and a penalty parameter for the error term of 10^*-*3^. We used 10-fold cross-validation. To produce the presented data, we used 9 out of 10 groups for training and one (i.e. 10% of data) for testing. This process was repeated 10 times so that each group was used for validation only once. In order to verify that our system performance was not simply due to overfitting of the regression method for the set of durations we included, rather than the ability of the system to estimate time, we tested the model estimation performance when excluding some durations from the training set, but keeping them in the testing set. The mean normalised error for durations included and excluded in each experiment is shown in (Fig. 9). As can be seen, only when excluding a large number of training levels (e.g. 10 out of 13 possible levels) does the estimation error get notably larger, suggesting that model performance is not attributable only to overfitting in the regression-duration estimates are robust across the tested range.

**Figure 9:**
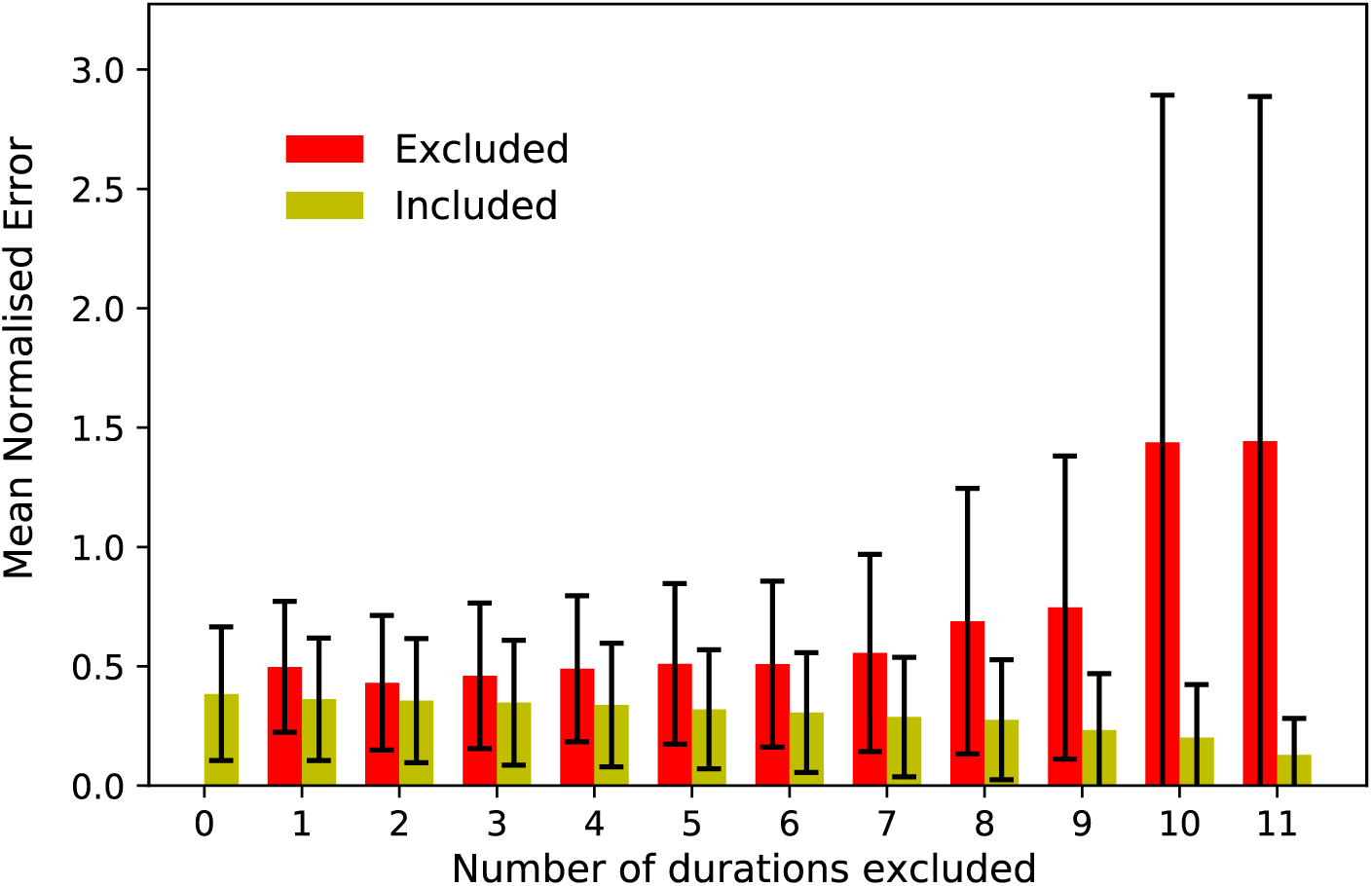
**(Supplementary figure)** Comparison of system performance by means of normalized duration estimation error, when a subset of testing durations were not used in the training process. For each pair of bars, 10 trials of N randomly chosen durations (out of 13 possible durations) have been excluded (x-axis). The support vector regression was trained on the remainder of the durations and tested on all durations. The errors for excluded and included trials are reported for each N. Only when excluding a large number of training levels (e.g. 10 out of 13 possible levels) does the estimation error get notably larger.

### Changes in classification network activation, not just stimulation, are critical to human-like time estimation

As outlined in the Introduction, our proposal is built on the idea that changes in the sensory environment, as reflected by neural activation within sensory processing networks, provide a mechanistic basis for human time perception. In a minimal interpretation, one might suspect that the efficacy of our model (including the basic ability of the model to estimate time, that model estimates improve with human-like gaze constraints, and that estimates are biased in different scenes in a way that follows human reports) may reflect only the basic stimulus properties. This interpretation would mean that the use of a human-like sensory classification network adds little to our understanding of duration estimation generally, or more precisely, the role of sensory classification networks in human time perception. To examine this issue we conducted a series of experiments wherein, rather than using the difference in network activation to indicate salient difference, we directly measured the Euclidean distance, by pixel, between successive frames of the stimulus videos. As in the initial experiments reported in the main text, we conducted these experiments under two conditions: one condition in which each frame of the video was constrained by human gaze data (“Gaze”), and another condition in which the whole video frame was used (“Full-frame”). In both cases, as with the initial experiments, the difference between successive frames was compared to a dynamic threshold, detected salient differences accumulated during a test epoch, and support vector regression trained on the accumulated salient differences and the physical labels of the interval in order to produce estimates of duration in seconds (as detailed in the methods for the main model). Consequently, any potential difference in results between these experiments and the experiments reported in the main text, conducted based on activations within the classification network, indicate the contribution of the perceptual processing within the classification network to time perception.

As can be seen in Fig. 11, estimates of duration can still be produced based on the pixel-wise differences in the video for both “Gaze” constrained video, as well as for the “Full-frame” video, as indicated by non-zero slopes in estimation. This basic sensitivity to duration is not surprising, given that our model of time perception is based on perceptual changes driven by sensory signals. Crucially, though, these results show several clear differences to both the classification network-based estimates, as well as human reports. Most obviously, estimation when using the “Full-frame” video is much poorer than for either of the “Gaze” or “Full-frame” models reported in the main text, with short durations dramatically overestimated, and estimations for office and cafe scenes similarly underestimated. These findings are clearly reflected in the mean deviation of estimation, shown in Fig. 10. While the overall pattern of biases by scene for the “Full-frame” video replicate the same pattern as for human reports (city *>* campus/outside *>* cafe/office; see Fig. 3G in the main text), both the overestimation of scenes containing more change (city and campus/outside) and the underestimation of the scenes containing less change (office/cafe) are much more severe. Overall, poor estimation performance when using “Full-frame” video is attributable to the estimation being driven only by the pixel-wise changes in the scene, especially for scenes wherein very little changes between successive frames on a pixel-wise basis (office/cafe; green line in Fig. 11). In these scenes, there are many instances where the scene remains unchanged for extended periods, therefore producing no pixel-wise differences at all with which to drive estimation.

**Figure 10:**
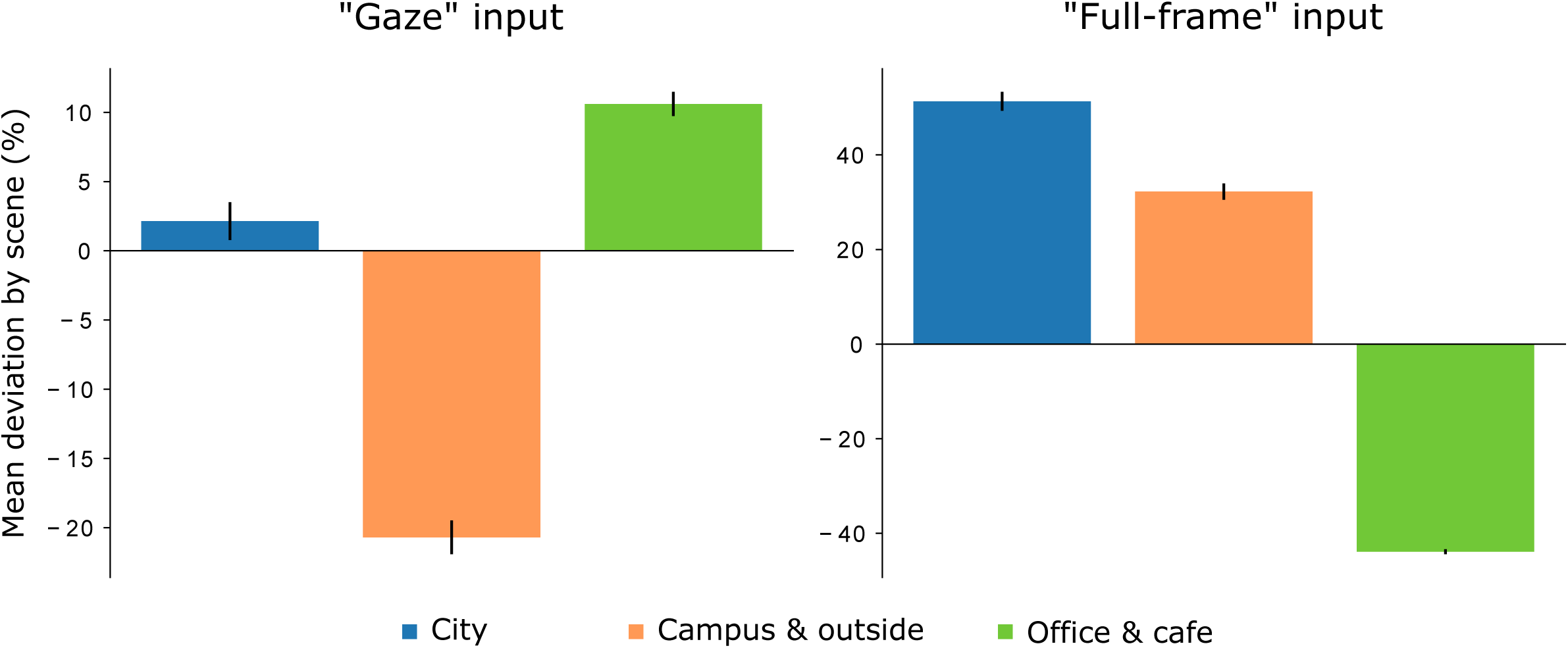
**(Supplementary figure)** Mean deviation of duration estimations relative to mean duration estimation, by scene type. Estimations were produced based on the raw Euclidean distance between video frames, by pixel, rather than using classification network activation. Left panel shows deviation of estimations, based on videos constrained by human gaze (“Gaze” input; as in human and model experiments in the main text), the right panel shows the same based on the “Full-frame” of the video. Error bars indicate standard error of the mean.

**Figure 11:**
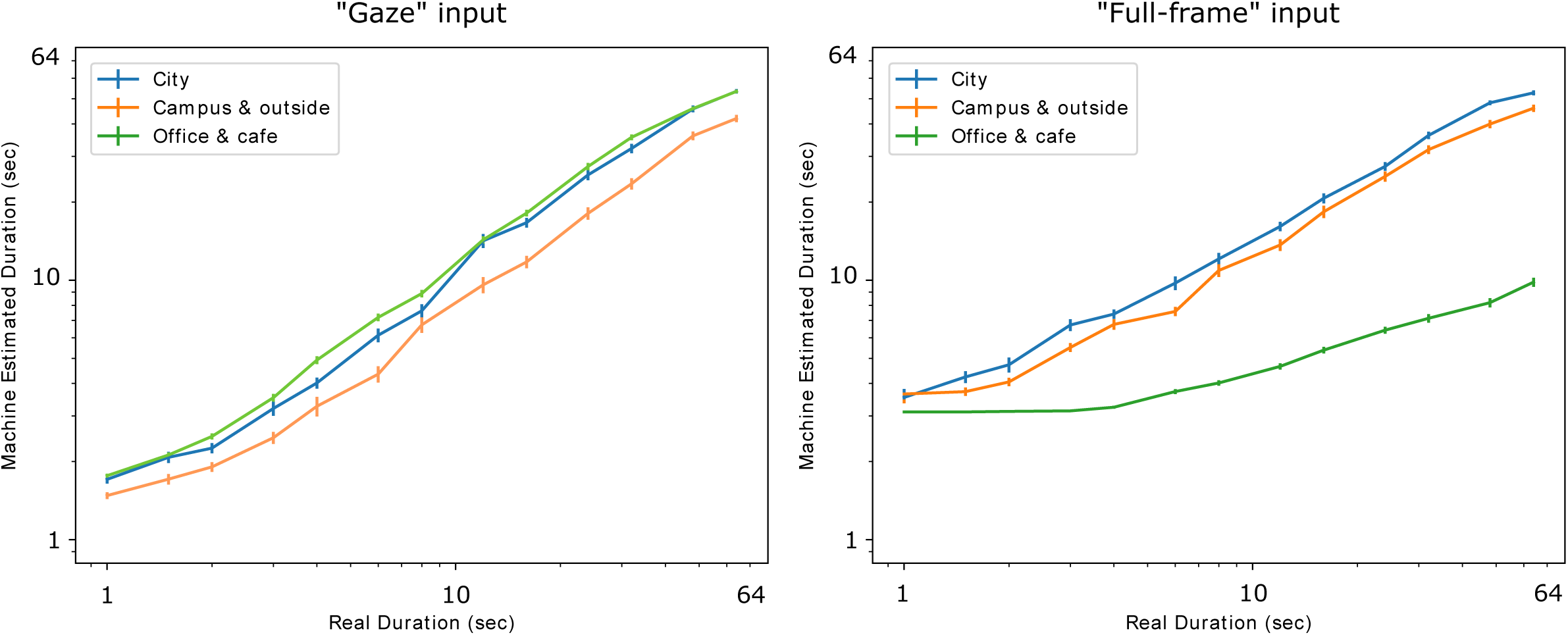
**(Supplementary figure)** Duration estimation for the 13 tested video durations, by scene-type. Estimations were produced based on the raw Euclidean distance between video frames, by pixel, rather than using classification network activation. Left panel shows estimations based on videos constrained by human gaze (“Gaze” input; as in human and model experiments in the main text), the right panel shows estimations based on the “Full-frame” of the video. Error bars indicate standard error of the mean.

By contrast, estimations based on gaze-constrained video (“Gaze” input) show superior performance to those for the “Full-frame” input video, with better slope of estimation (Fig. 11), and less severe over/underestimation. These results support the findings reported in the main text regarding the importance of where humans look in a scene to the estimation of duration. However, as is clearly depicted in Fig. 10, when considering the pattern of biases induced by different scenes, estimations based only on gaze-constrained video do not replicate the pattern of results seen for both the classification network-based model and human estimations (Fig. 3G and H) reported in the main text. Rather, estimations based on gaze-constrained video alone substantially underestimate durations for scenes based in the campus/outside, while overestimating the scenes with the least perceptual change (office/cafe scenes).

Overall, these results show, consistent with the proposal outlined in the Introduction, that the basis for human-like time perception can be simply located within changes in sensory stimuli. More importantly, they also show that it is not just the sensory stimuli alone that drive time perception, but also how stimulation is interpreted within perceptual classification networks. By basing our core model, as reported in the main text, on stimulus-driven activation in a human-like visual classification network, our model is able to naturally capture human-like biases in duration estimation in a way that is not possible based on the sensory stimuli alone.

## Notes

1 Note that relative over and underestimation will partly depend on the content of the training set. In the described experiments, the ratio of scenes containing relatively more changes, such as city and campus or outside scenes was balanced with scenes containing less change, such as office and cafe. Different ratios of training scenes would change the precise over/underestimation, though this is similarly true of human estimation, as the content of previous experience alters subsequent judgements in a number of cases, e.g.^36,54,55^. See Methods for more details of training and trial composition.

2 Presumably humans don’t experience time only in seconds. Indeed it has been shown that even when they can’t provide a label in seconds, such as in early development, humans can still report about time ^68,69^. Learning the mapping between a sensation of time and the associated label in standard units can be considered as a regression problem that is solved during development ^70,68^

